# Computational Justice: Simulating Structural Bias and Interventions

**DOI:** 10.1101/776211

**Authors:** Ida Momennejad, Stacey Sinclair, Mina Cikara

**Affiliations:** Columbia University; Princeton University; Harvard University

## Abstract

Gender inequality has been documented across a variety of high-prestige professions. Both structural bias (e.g., lack of proportionate representation) and interpersonal bias (e.g., sexism, discrimination) generate costs to underrepresented minorities. How can we estimate these costs and what interventions are most effective for reducing them? We used agent-based simulations, removing gender differences in interpersonal bias to isolate and quantify the impact and costs of structural bias (unequal gender ratios) on individuals and institutions. We compared the long-term impact of bias-confrontation strategies. Unequal gender ratios led to higher costs for female agents and institutions and increased sexism among male agents. Confronting interpersonal bias by targets and allies attenuated the impact of structural bias. However, bias persisted even after a structural intervention to suddenly make previously unequal institutions equal (50% women) unless the probability of interpersonal bias-confrontation was further increased among targets and allies. This computational approach allows for comparison of various policies to attenuate structural equality, and informs the design of new experiments to estimate parameters for more accurate predictions.

---

Women and minorities remain underrepresented in the highest ranks of STEM fields (*1*), politics (*2*), business (*3*), and other high-prestige professions (*4*). This underrepresentation at higher ranks exists despite more representative demographic distributions in early-career stages (e.g., in STEM, (*1*, *5*, *6*)). Losing the talents of a wide swath of the work force incurs tremendous costs both to individual workers’ earning potential (*7*) and to institutions’ earnings, innovation, and decision-making (*8*, *9*). That said, there are costs to persisting in fields in which one represents a minority. For example, pervasive racial discrimination in industrialized countries such as the U.S., U.K., and Australia incur measurable mental health costs for persons of color in the work force (7, 8). These cumulative costs are a significant driver of organizational exit for women and other minorities (*10, 11*).

Most interventions designed to remedy underrepresentation focus on changing individual factors, such as personal attitudes or stereotypes, or individuals’ interpersonal behaviors, such as reducing bias in hiring decisions (*12*). However, evidence indicates that ‘diversity’ trainings directed at these factors yield mixed results at best (*12*, *13*). Moreover, they tend to focus on recruitment rather than retention.

Here we propose that a key contributor to underrepresentation is the increased costs associated with being underrepresented in the first place. In other words, structural disparities (e.g., a historically male-skewed gender ratio) generate conditions under which bias will accumulate in greater proportion toward minorities (e.g., women) who are present to receive it. Thus, systemic or structural interventions, though less explored in professional settings, hold promise as key to institutional change, e.g., quotas in politics (*14*). Despite the efficacy of structural interventions, there are prohibitive factors that leave them less explored; for example, structural interventions are logistically challenging, are expected to show effects only over long time scales, and carry potential risk of “over-correction” or failure. As such, many organizations and institutions may consider it too risky to test the effects of large-scale structural change. Even for institutions willing to implement structural change policies, empirical parameters are lacking to estimate over which time horizons different strategies for structural change would yield parity (e.g., 1 year, 10 years, 50 years?).

The present work attempts to overcome challenges associated with testing the effects of structural disparity, and the long-term efficacy of structural and interpersonal change policies. To do so, we conduct multi-agent simulations (*15*–*18*), parameterized by empirical evidence from gender and racial bias research, and compare the accumulated costs of disparity under different gender ratios and intervention strategies. Specifically, we compare the long-term efficacy of objection to bias (interpersonal) and an equality policy (structural) for reducing the costs of gender disparity to individuals and institutions. Similar approaches have successfully assessed the emergence of segregation (*15*) and long-term effects of racial profiling policies (*16*). For simplicity, here we focus on gender disparity and leave simulating racial disparity and intersectionality to future work.

## Overview

Here agents interact in simulated meetings. In each meeting, one or more agents can make sexist comments with varying probabilities and, when they do, other agents can object to the occurrence of sexism with empirically derived probabilities. Furthermore, in each meeting, agents learn from the observation of sexist comments and corresponding objections: the probability of sexism either increases, when sexism occurs with no objection, or decreases when objections follow. These learning parameters are set based on empirical results (*19*).

Our simulations quantify (i) the impact of unequal gender ratios on the count of sexist encounters, (ii) the associated costs of gender bias to individuals and institutions, and (iii) the long-term impact of equality policy and bias confrontation. We based the multi-agent simulation on the Petrie multiplier, a thought experiment proposed by Karen Petrie, and added multi-agent learning from observation, objection probabilities, and costs—of receiving sexist comments, objecting to sexism, and receiving objections. To maximize the relevance of our simulation results to real world solutions, all parameters were estimated following our literature review of human behavioral data. When there were no studies from which to directly estimate the parameters, we derived simulation parameters from related empirical evidence (see Methods).

## Methods

We simulated institutions, each initialized with 100 agents with a given gender ratio (e.g., 20 female), parameters for the probabilities of agents displaying and objecting to bias, and parameters for observational learning (contagion). We then simulated 1000 sequential *meetings*, in each of which 2-8 agents were randomly sampled from the population to interact (*20*). In these meetings simple rules determined whether bias occurred, whether it was objected to, and how all this affected costs and future probabilities of bias (see below). We repeated this simulation 1000 times and computed changes in probabilities and costs (Fig 1). It is important to note that we initialized every simulation with the assumption that male and female agents had equal probability distributions for displaying anti-female and anti-male bias, respectively, as well as equal costs and learning rates for changes in their bias probabilities. These conservative assumptions ensured that any disparity we observe in the overall costs and in the growth of bias are due only to gender ratio.

**Figure 1.**
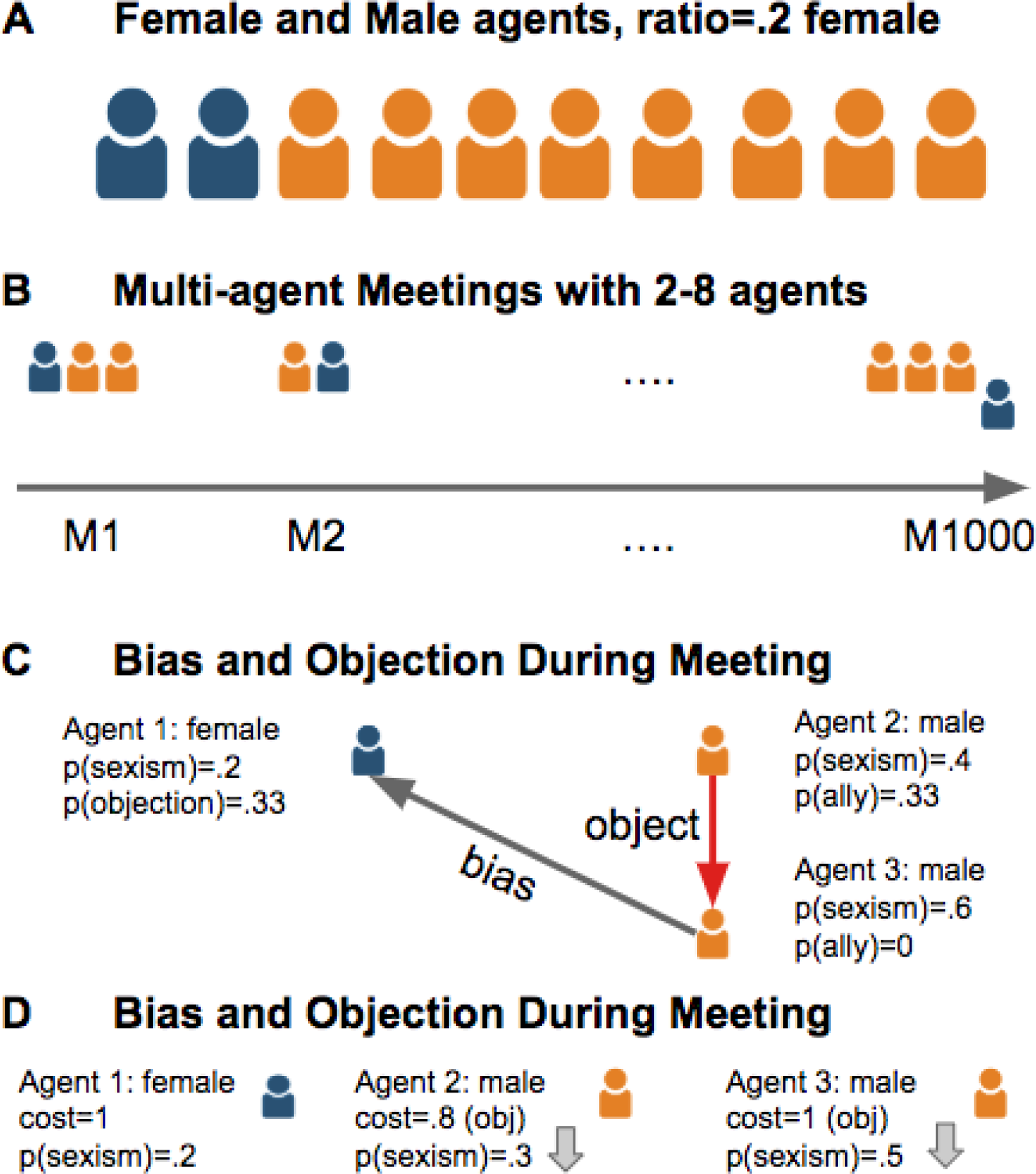
Agent based simulation. (A) We simulated each institution with 100 agents with different gender ratios. For instance, in a .2 female institution 80% of agents were male and 20% female. The agents were drawn from populations with equal probability distributions for sexism. That is, if the institution’s ratio was .5, the mean and standard deviation of their sexism probabilities were equal. (B) In each institution we simulated 1000 meetings (*M*) with 2-8 agents randomly drawn from the pool of 100. Each simulation with gender ratio and confrontation probabilities was repeated 1000 times. (C) During each meeting agents interacted. With some probability (set based on empirical findings) agents made biased comments, leading to a cost of 1 to the target. Agents from the target group (e.g., female agents) or allies (e.g., male agents) could object. If an objection was made, the perpetrator received a cost of 1. Making an objection also incurred a cost of 1 when made by a target and a cost of .8 when made by an ally. (D) When bias occurred without objection, agents from perpetrator group increased their sexism probabilities by .1. When bias was followed by an objection, the perpetrator group decreased their sexism probabilities: by .1 if an in-group made the objection, and by .05 if an agent from the other group (i.e., an ally) objected.

### Probabilities of expressing bias, or sexism probabilities

The initial probability distribution for giving a sexist comment in a meeting was set equal for male and female agents. In absence of empirical estimates and inspired by other implementations of the Petri multiplier (*20*–*22*), we initialized parameters such that 50% of agents of either gender had a 0 probability of making a sexist comment, 10% had a probability of .2, 10% probability of .4, 10% probability of .6, 10% probability of .8, and 10% probability of 1. An agent could make a sexist comment during a meeting if either (a) there were equal numbers of male and female agents, e.g., 2 male and 2 female, (b) or if the agent’s group outnumbered the other, e.g., 3 males and 1 female, and (c) a random number, generated during each meeting, was smaller than its probability of objection (e.g., coin flip gives .7, the agent will make a biased comment if its bias probability is >.7). Giving a biased comment had no costs in itself (unless it was met by objection, see below). A biased comment targets all agents of the other gender in the meeting. After the biased comment, all agents in the meeting with the *target* gender receive a cost of 1. Costs are additive (e.g., being the target of 5 agents leads to a cost of 5).

### The probability and cost of objecting to sexism towards one’s own group

When bias occurred during a meeting, an agent in the meeting whose group was *targeted* with bias could confront the bias. The distribution of probabilities for objecting as target were as follows: of all agents of each gender 20% object to bias with .33 probability, 10% object with .66 probability, and 10% with probability 1. This is based on empirical studies in which 35% to 45% of women objected to sexist encounters (*23*, *24*), and of this number half responded to 1 out of 3 sexist comments, 10% to 2 out of 3, and 10% to all 3 sexism comments (*23*). In the absence of similar studies for men, we made the conservative assumption that the same distribution is true for male agents here. There are well-documented career and social costs when an agent objects; these stakes are even higher in professional settings and may inhibit objection to bias (*25*). As such, we generated a penalty cost of 1 when an agent objected on behalf of his or her own *target* group. (Future versions of the model can accommodate different costs, e.g., depending on differences in ranks, the social influence of an agent, and stakes of the meeting.)

### The probability and cost of objecting as an ally

When bias occurred in a meeting, both target and *non-target* agents in the meeting (i.e., agents with the same gender as the perpetrator) could confront. This is bias confrontation as an *ally*. While we do not have direct empirical studies on the probability of objecting as an ally, we estimated ally probability based on a study of perception of sexism (*26*)(see Supplemental Material). To determine costs, we relied on empirical evidence indicating that men who confront sexism against women are seem about 10% more likeable than women who confront sexism against women (*27*). Furthermore, studies show a stronger preference for social exchange with third-party observers who punish unfair treatment but not victims who punish (*28*), perhaps since third-party punishment by men appears to be against self-interest and hence more morally driven (*29*). Based on these findings, and in absence of direct empirical evidence, we set the cost for objecting as an ally to .8 (20% less costly than objecting as a target).

### Learning from bias and objection

Studies on social learning by observation of bias and bias confrontation are rare. We set our agents’ learning parameters based on a study examining the effect of bias confrontation on Twitter (*19*). Twitter users who used racial slurs reduced their probability of using slurs by about 10% when an in-group member, i.e., a white male, confronted them and by about 5% if confronted by an out-group member, i.e., a black male. It is worth noting that these numbers are averaged across effects of twitter users with high and low followers.

In our model when sexism occurred in a meeting (of 2-8 agents) and there were no objections, non-target agents (same gender as perpetrator) increased their probabililty of sexism by 10% of their original probability. For instance, if a male agent’s bias probability was .2, after a sexist comment by another male agent occurred with no objection, the observer’s probability increased to .22. When a biased comment during a meeting was objected to by an ally, then all non-target agents in the meeting reduced their sexism by 10% of their original probabilities (e.g., .2 decreased to .18). When the objection was made by an agent from the *target* gender, then the probability of future sexism by *non-targets* was only reduced by 5% of its original, (e.g., .2 decreased to .19).

Note that agents’ probabilities of making biased comments in future meetings followed from the updated probability. In short, our agents *learned* from the occurrence of bias and confrontation, such that their probability of future bias could increase (bias propagation) or decrease (bias inhibition).

## Results

### Gender ratio affected the frequency of biased interactions

We first tested the effect of gender ratios in simulated institutions on the mean count of anti-male and anti-female comments per meeting (Figure 2). Sexism probabilities were initialized as equal in all simulations with equal distributions of gender bias and bias confrontation for male and female agents. As such, in an institution with a 50/50 gender ratio the count of anti-female and anti-male comments were equal (Figure 2, permutation test’s *p*=.*9221*), while in an environment with gender ratios of .1, .2, .3, and .4 there were much higher anti-female than anti-male comment counts (Figure 2, permutation test’s *p*<10^−5^). These effects were exacerbated in absence of objections to interpersonal bias (Supplementary Figure 1).

**Figure 2.**
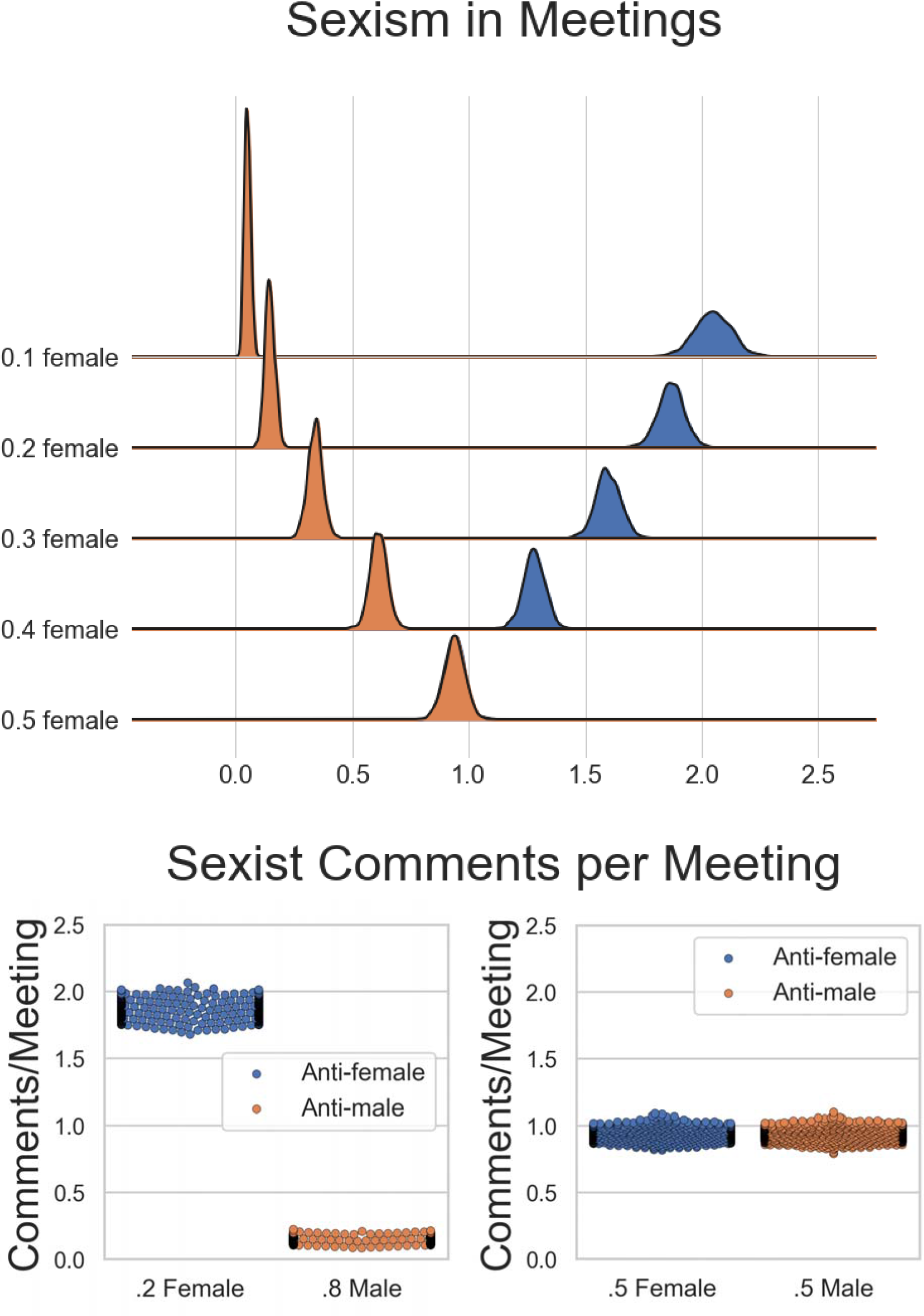
Mean number of sexist comments per meeting. (Top) Sexist remarks received by female agents (blue) and male agents (orange) per meeting plotted over 1000 meetings in simulated institutions initialized with different gender ratios (.1 female, .2 female, .3 female, .4 female, .5 female) and equal sexism probability distributions for agents of both genders. The Y axis of distribution plots, for each institution with a given ratio, indicate the number of agents receiving the corresponding number of comments on the X axis. (Bottom) Swarm plots of distributions from the plot above for .2 female ratio and .5 female ratio for comparison. The Y axis indicates mean comments per meeting in an institution and each dot represents the comments received by women (blue) or men (orange) in one of the 1000 simulations of a given institution.

### Probabilities of bias increased over time

We compared the post-simulation probability distributions for expressing bias (sexism probabilities) after 1000 meetings in environments across different gender ratios (.1, .2, .3, .4, or .5 female). Again, initial bias probabilities were equal for male and female populations in the initialization of an environment. Notably, we allowed observational learning in accordance with empirical findings (*19*): sexism probabilities could increase due to observing sexism that received no objection, or decrease due to observing objection. We observed that gender ratio disparity affected the increase in sexism probabilities (Figure 3). Probabilities of expressing bias grew more in the majority agents (e.g., male) in institutions with low ratios of female agents (Figure 3, right), whereas these probabilities and their growth were equal in 50/50 institutions (Figure 3, right).

**Figure 3.**
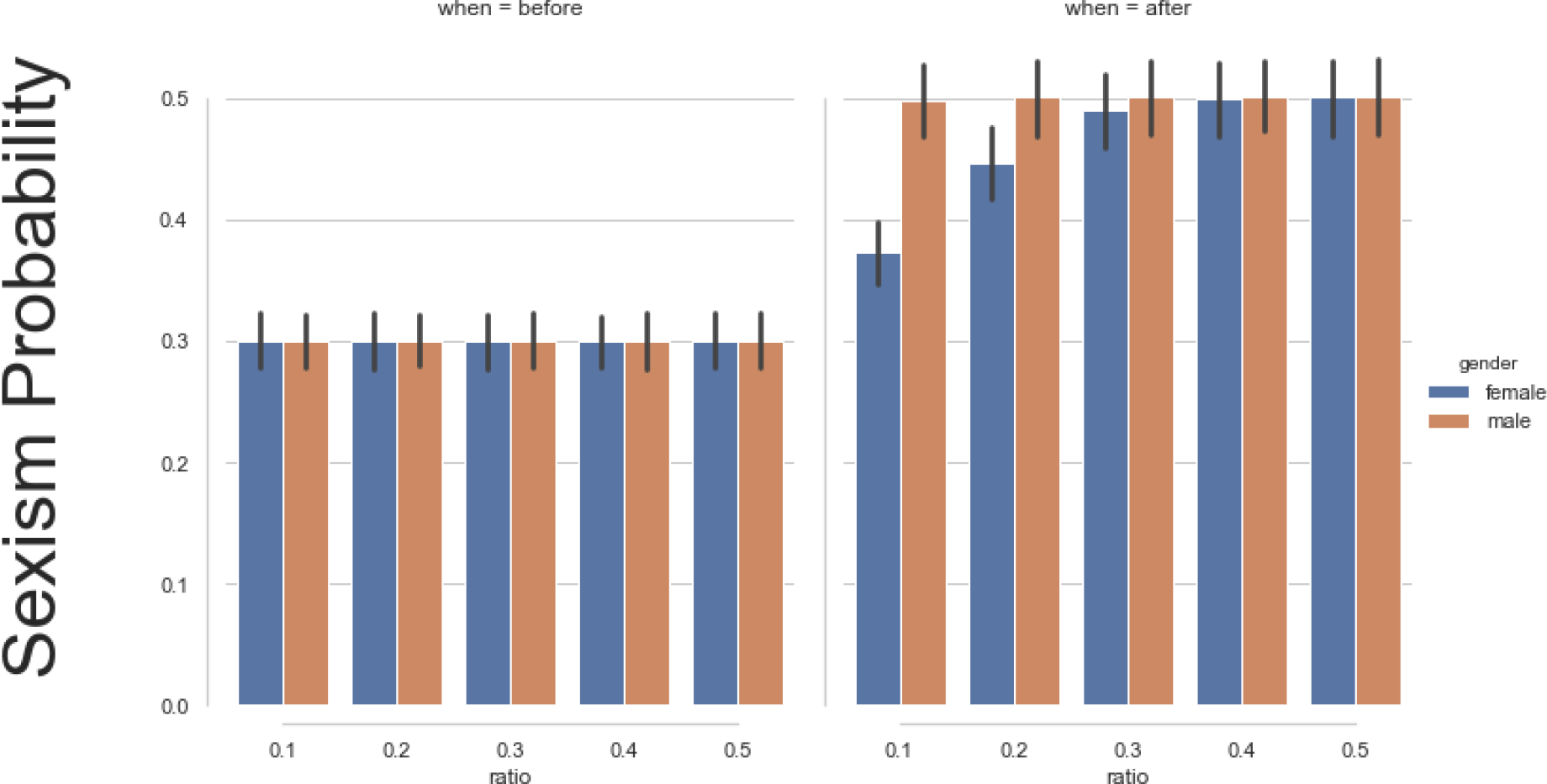
Mean probabilities of interpersonal bias before and after 1000 meetings. (Left) Mean interpersonal bias probabilities before 1000 meetings in institutions initialized with different gender ratios (.1, .2, .3, .4, .5 female). (Right) Mean interpersonal bias probabilities after 1000 meetings in institutions initialized with different gender ratio. (Male: orange, Female: blue).

### Cumulative costs of bias to individuals and the institution

There are three kinds of costs in the present model, inspired by empirical findings (*24*, *25*, *27*): the cost of being the *target* of bias (target agents receive cost of 1), the cost of *confronting* bias (target objector receives cost of 1, ally objector receives cost of .8), and the cost of *being* confronted (perpetrator receives cost of 1). For each simulated institution with a given gender ratio (e.g., .2 or .5 female) we ran 1000 simulations using these empirically set parameters and estimated the total cumulative costs of bias and confrontation for female and male agents (Figure 4). Furthermore, to compare the total cost of bias to institutions with different gender ratios, we summed the costs of bias to female and male agents for each environment (Figure 5). Moreover, the absence of objections did not reduce the costs to institutions (Supplementary Figure 2) but significantly worsened the costs to women (Supplementary Figure 1). The more skewed the gender ratio, the higher the costs to women and to institutions as a whole. More balanced gender ratios yielded a large decrease in costs to women and institutions, while moderately increasing the costs for men (Figure 4).

**Figure 4.**
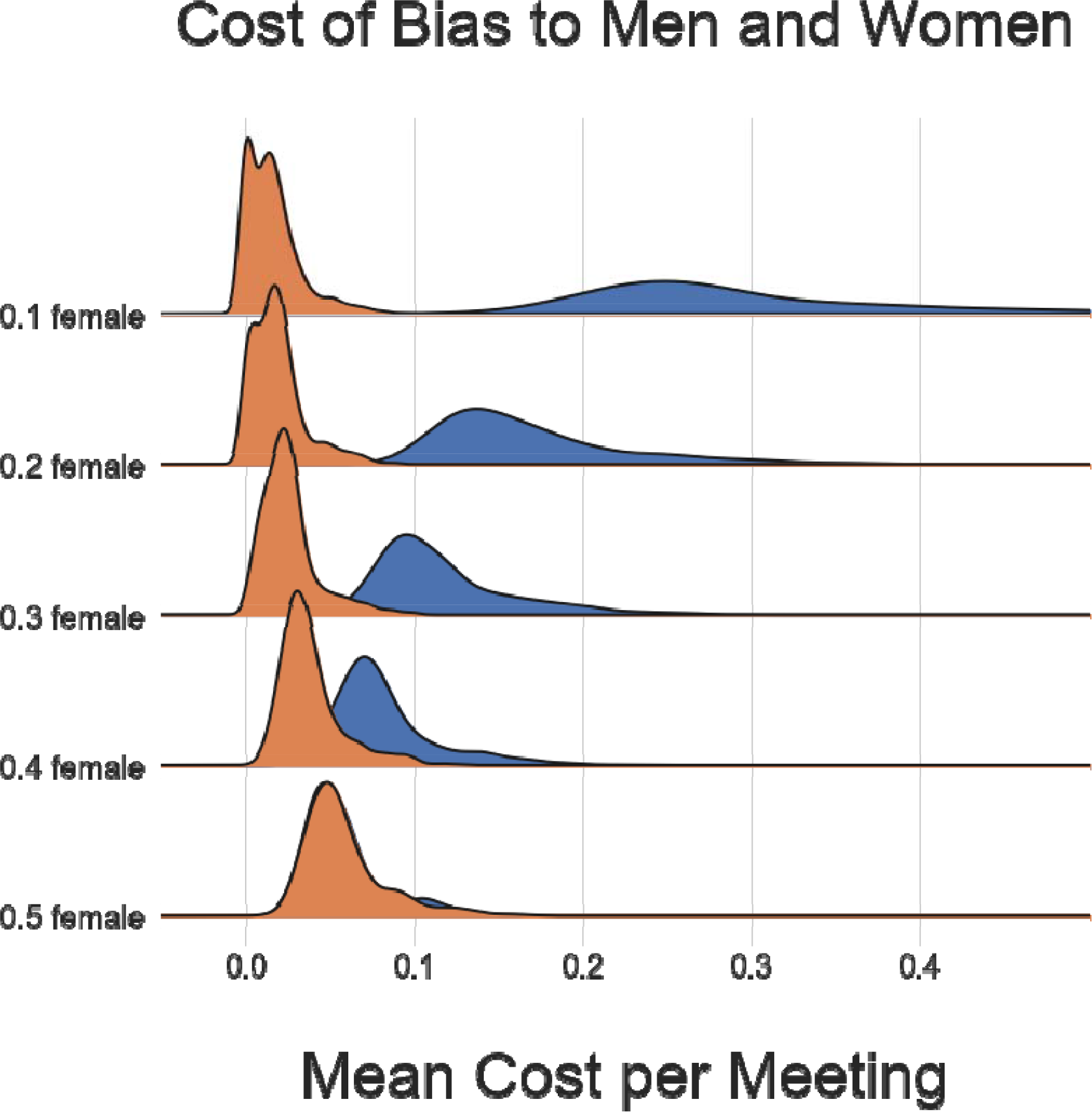
The cost of bias for male and female agents. Total costs for female agents (blue) and male agents (orange) arrived at after 1000 simulations of 1000 meetings initialized with different gender ratios (.1 female, .2 female, .3 female, .4 female, .5 female). The Y axis of the distribution plots, for each institution with a given ratio, indicates the number of agents receiving the corresponding costs on the X axis.

**Figure 5.**
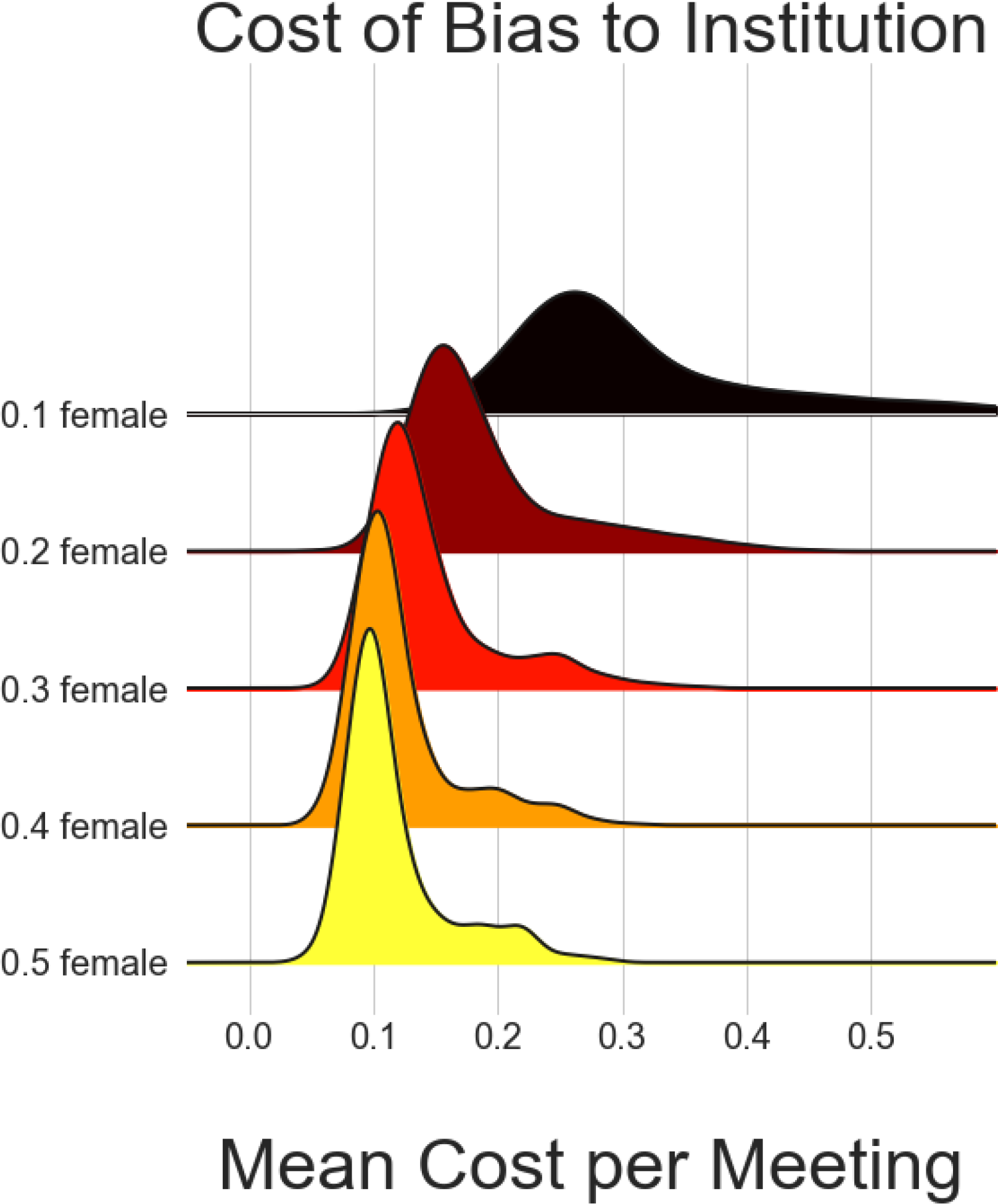
Cost of bias to institutions. Cumulative costs of bias for institution arrived at after 1000 simulations of 1000 meetings initialized with different gender ratios (.1 female, .2 female, .3 female, .4 female, .5 female).

### Simulating the introduction of an equality policy

Here we tested whether enforcing gender parity in an institution with a previously skewed gender ratio would eliminate sexism. Specifically, after 1000 meetings with a 20/80 ratio we simulated enforcing a 50/50 equality policy. We restarted the environment with a .5 ratio but carried over the probability of sexism arrived at after the 1000 meetings in the .2 environment. We found that the number of sexist comments per meeting (Figure 6) and the costs to men and women (Supplementary Figure 3) were far from equal, and the gap widened with further meetings.

**Figure 6.**
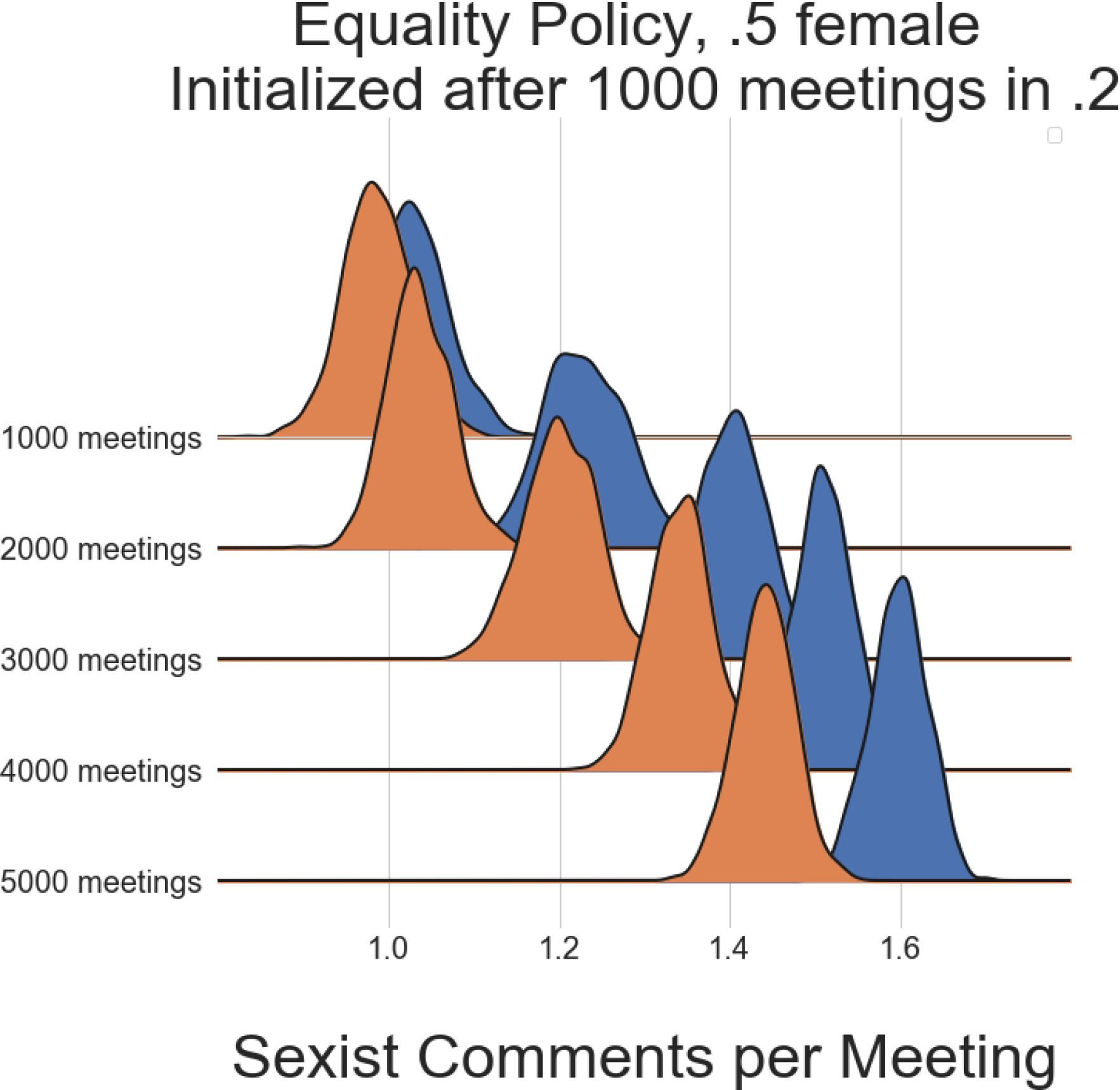
Enforcing equality policy does not yield equal sexist encounters. We computed sexism probabilities for male and female agents after 1000 meetings in a .2 environment, and initialized a new environment with .5 ratio carrying over the previous environment’s updated probabilities. We simulated 1000, 2000, 3000, 4000, and 5000 meetings in the new 50/50 environment to compare the long-term effects of this equality policy. In spite of the 50/50 gender ratio in the new environment, we observed a large gender gap between sexism comments received by male and female agents per meeting which increased over time.

### Combined effect of equality policy and (increased) objection

Here we tested whether instituting an equality policy plus (increased) objection would eliminate gender differences in the receipt of sexist comments (Figure 7). Only when we increased the objection probabilities such that 20% more agents from the population—10% targets and 10% more allies—would object with probability = 1 did the simulated institutions overcome the gender gap in receipt of sexist comments.

**Figure 7.**
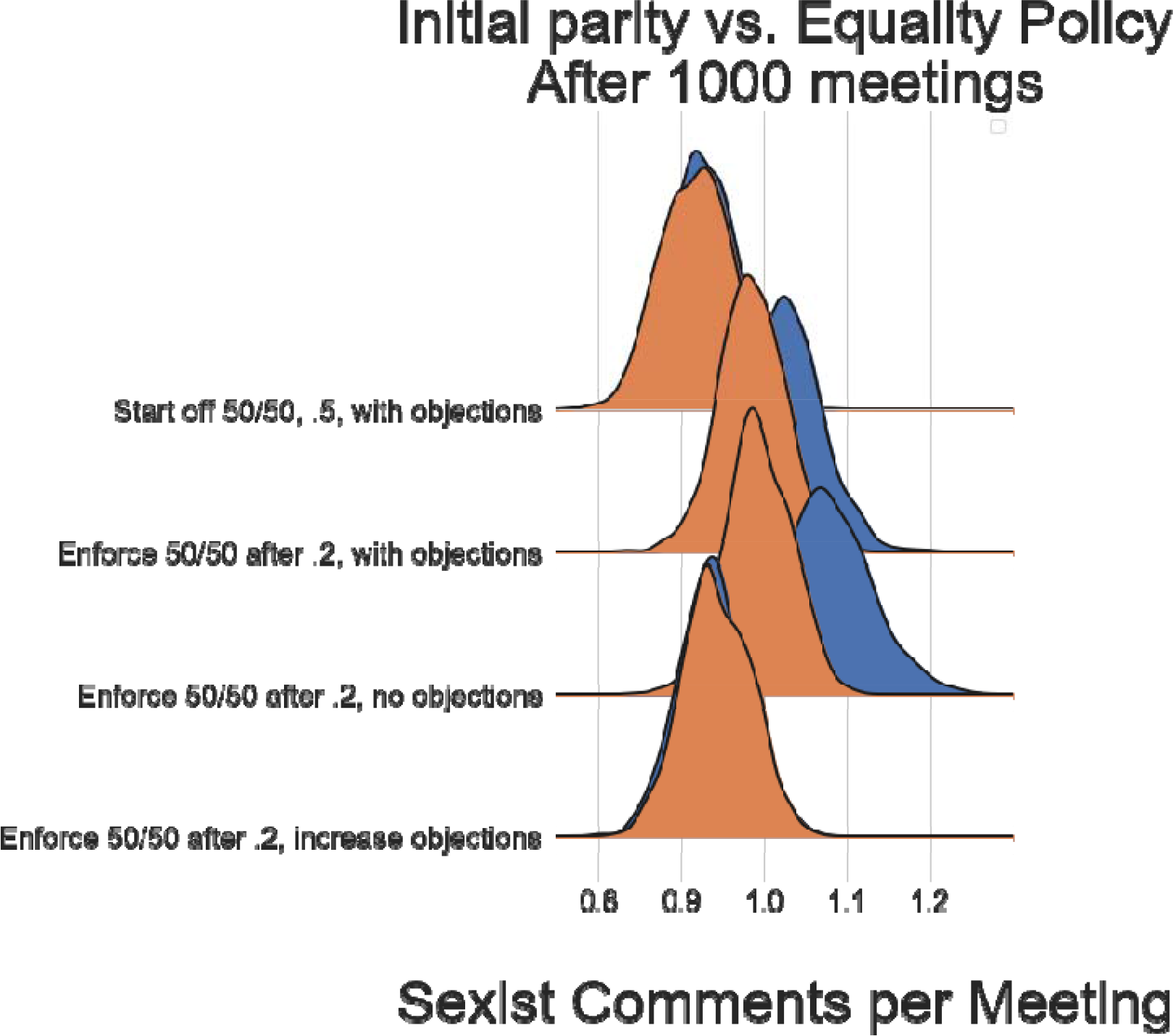
Initial parity and equality policy with varying objections. We compared sexist comments per meeting over 1000 simulated meetings in four institutions. The first institution was initiated with a 50:50 gender ratio and equal probabilities of sexism for men and women. Objections were allowed. Here the number of sexist comments were equal. The second institution was initiated with a .5 ratio but carrying over sexism probabilities derived after 1000 meetings with a .2 ratio. We refer to this as enforcing an *equality policy* to a previously .2 environment. The third institution was the same as the second, but without allowing objections by either targets or allies. Unsurprisingly, we observed that removing objections increased the gender gap in the number of sexist comments, but this allowed us to isolate the relative efficacy of objections. Critically, the fourth institution was initiated the same as the second and third, but we increased the objection probabilities such that 20% more agents from population would object to sexist comments with probability = 1. In this environment, the gender gap in the number of comments disappeared. In all simulations, objections to sexism were required to avoid conditions worsening over time.

## Discussion

Here we simulated structural gender inequality and interventions for social change. We modeled interpersonal bias, due to individual differences in bias probability, as well as structural bias, due to unequal gender ratios. We ran agent-based simulations, excluding any gender differences in the probability of exhibiting interpersonal bias to isolate the effect of structural bias on our outcomes. This allowed us to quantify the costs of structural bias to individuals and institutions and compare the efficacy of interventions for attenuating said costs. We found that simulating institutions with unequal gender ratios (i.e., female minority environments) led to increased bias towards and costs for the minority group. Furthermore, confronting interpersonal gender bias by the target group and allies attenuated, but did not eliminate, the growth of bias and the costs associated with structural bias (i.e., skewed gender ratios). Notably, institutions with higher gaps in gender ratios suffered more costs (Figure 5).

We also tested the effects of an *equality policy*, enforcing a 50/50 gender ratio in a previously gender-skewed institution (20/80). After 1000 meetings, we found that the costs to male and female agents were far from equal and this gap widened with more meetings (Figure 6). Thus, enforcing an equal gender ratio alone will not eliminate bias from institutions with a history of structural disparity. However, increasing the participation of allies in objecting to sexism could (Figure 7). Thus a combination of both structural and interpersonal policies for social change appears to be necessary for promoting equal treatment of men and women in professional settings.

Note, however, that there are many ways in which our simulations represent a ‘best case scenario.’ For example, we assumed men and women have equal probabilities of displaying bias, and are equally likely to confront when they detect bias. Our simulations focused on binary groups for simplicity, and did not take into account the mediating effects of race and status, or intersectionality (*30*). Furthermore, in the present model an agent’s parameters only change during inter-group interactions. However, in the real world unjust gender norms can be learned during single-gender interactions (e.g., “locker room” talk where men enforce sexist norms, minority “gate keepers” who may impede other minorities to remain the exception, women showing gender bias against other women, etc.). We also did not model institutional exit—when agents opt to leave an organization because the cumulative costs of staying become too high (*11*)—because we did not have any data on which to base that threshold. In the real world, women and minorities who ascend the ranks become relatively less well represented over time as other women and minorities exit. Our stimulations suggest those who remain only experience an *increased* concentration of bias over time, which could account for the leaky pipeline. Even if individuals persist past early stages, the costs will increase over time until they eventually exceed an individual’s threshold.

More generally, this exercise serves to highlight several lacuna in the empirical literature. The main challenge we faced in specifying these models was that we could not find studies that directly tested the particular parameters we required for these simulations. For example, in these models we assumed men and women were equally harmed by sexist comments, equally punished for confronting sexism, and equally likely to confront on behalf of male and female targets, because we could not find evidence to suggest that we should assume otherwise. While several studies have highlighted that non-target confronters (e.g., men confronting sexism against women) are taken more seriously and seen as more legitimate (*27*, *29*), there are no data of which we are aware to speak to base rates in non-target confrontation, changes in incidence of bias (i.e., learning) following confrontation or its absence, and so on. Therefore this work highlights several future research directions that would help inform the generation of more realistic models, and in so doing, the development of more effective strategies for bias reduction.

### Limitations and future directions

In order to simplify this first attempt at modeling bias interventions, we restricted our simulations to binary gender identities. However, future simulations should implement intersectionality—assigning multiple social group memberships to each agent—in a similar model.

As we note above we also did not allow agents to leave the institutions. In the real world, however, individuals drop out of institutions or professions due to the accumulating costs of being a minority. Findings from twenty corporations in 2008 indicated that women and minorities were more likely to quit jobs than their majority counterparts (*10*). Future simulations should incorporate this dropout threshold to account for the “leaky pipeline”, where gender inequality increases at the top. In this case, due to increasing accumulated costs the longer minorities remain in unbalanced institutions.

Future simulations could also attempt to capture the influence of norms (e.g., whether each agent *believes* that the majority or minority of agents in a given meeting are biased). In complement, future empirical work informed by these models can quantify the impact of explicitly stating anti-bias norms and instituting public disciplinary objections on bias propagation and inhibition. Furthermore, recent studies suggest that third-party punishment combined with compensation to victims of unfair treatment may have a stronger effect than mere punishment (*28*). Future studies can simulate the long-term effects of policies that combine stronger third-party or ally interventions and the compensation of targets proportional to the unfairness they have experienced.

Finally future simulations should accommodate the nested and hierarchical network structure of these interactions and how they are influenced by and impact ties between agents, which may be crucial in career advancement. For example, institutions can be modeled as networks of social interactions with specific graph structures. It is then possible to quantify the graph properties of each agent as a node in a broader network, e.g., eigenvector centrality, and measure the efficacy of their objections depending on their network influence (*19*, *31*, *32*) as well as changes in their network ties as a function of social learning, e.g., reducing ties to perpetrators or objectors with different effects on structural change (see (*33*) for similar approach to increasing efficacy of anti-bullying campaigns to change school norms and students’ behavior)

An important contribution of the present approach is that it can inform the design of new experiments and the acquisition of missing data. The exercise of generating the necessary parameters for agent-based simulations identifies precisely where data are lacking.

In conclusion, we think these simulations can be used to quantify the effects of inequality and test potential interventions in a risk-neutral fashion. In turn, simulations can guide future experiments and data collection efforts to inform realistic model parameters. Via this iterative process, empirically parameterized simulations can help identify effective policies for confronting and reducing bias. Collective efforts in machine learning, multi-agent simulation, data mining, and behavioral experiments will lead to more realistic simulations of how bias impedes career development as well as policies for social change. We believe this approach can be tailored to the need of small, national, and global institutions, and will offer improved strategies for institutions to achieve their stated goals and values.

## * Acknowledgement

We would like to thank Luis Piloto for insightful conversations and a Matlab implementation of earlier ideas, Vassiki Chauhan for helpful conversations, and Angie Michael Meiztler for organizing a workshop to teach the model to the public. We gratefully acknowledge the support of NSF CAREER award 1653188 (awarded to MC).

## Supplemental Material

Karen Petri’s thought experiment (*20*) suggests that in an environment with an 1/*r* female to male ratio (assuming equal distribution of sexism probabilities for men and women) women are *r*^2^ times more likely to receive a sexist comment. We designed a a multi-agent simulation inspired by this thought experiment, and extended the simulation by adding (a) social learning of bias probabilities, (b) probabilities of objection, and (c) costs of objecting and being objected to for sexism. All model parameters were derived from a review of empirical experiments on race and gender.

### Parameter selection and justification

#### Setting confrontation parameters

The literature on confronting sexism generally assumes that women are targets of sexism who may or may not chose to confront it when they experience it, and men are potential allies who may or may not chose to confront sexism against women. As such, there is no research examining the degree to which men confront sexism against men or women serve as allies for men who experience sexism. We did not build this assumption into the model. Because male and female agents in our model are initially equally likely to engage in sexist behavior, they both have the potential to experience being a target and an ally. For this reason, we used the empirical literature to determine the costs and likelihood of confronting bias against one’s own group and in the role of an ally (confronting bias against the other group), then applied these estimates equally to male and female agents depending on their role in the interaction.

#### Confronting bias against one’s group: Costs and likelihood

One strategy to mitigate gender bias would be for people to confront it every time they experience it. However, universally consistent confrontation does not occur because objection to injustice has personal and social costs that can impede one’s social repuptation, career, and health (e.g., Shelton & Stewart, 2004). Though confrontation can also serve to promote autonomy (Sanchez, Himmelstein, Young, Albuja, & Garcia, 2016). These costs include negative responses from others such as being perceived as less likeable, continued or increased harassment, losing a job or promotion opportunity, or receiving a poor grade.

Because of these costs, people do not confront biased behavior as often as they want or expect to. For example, 92% of women confront man asking sexually harassing questions when the cost is low (i.e., they have an alternative job offer), but only 22% confront when the costs are high (i.e., they are interviewing for their dream job; Shelton & Stewart, 2004).

Another study compared women’s tendency to confront perceived harassment in imagined versus real scenarios (Woodzicka & LaFrance, 2001). In the imagined scenario, 68% of women imagined they would refuse to answer sexually harassing questions during a job interview and 28% said they would either rudely confront the interviewer or leave the interview. However, during an actual face-to-face interview, 52% of women ignored sexist comments, only 36% asked polite clarification questions. Most important: <u>0% confronted the interviewer rudely.</u> The study concluded that faced with a harassing comment during a job interview, the fear of social rejection or jeopardizing a potential job changes the self-reported desire to launch assertive confrontation into, e.g., a polite question regarding the interviewer’s intentions. Finally, Swim and Hyers (1999) reported that 45% of women confronted a man who made a sexist remark (i.e., addressed at least 1 of 3 remarks) and only 16% did so directly (i.e. responded with comments such as “You can’t pick someone for that reason. Pick another person’’). Notably, the most frequent response to sexism was an expression of displeasure such as a sarcastic comment or an exclamation of surprise (Swim & Hyers, 1999). This 45% objection estimate replicated ten years later, when a survey study conducted by another group reported that only 46% of female undergraduates said they had confronted sexism in a past episode (Ayres, Friedman, & Leaper, 2009).

The breakdown of objections in the Swim and Hyers study informed our model parametrization. Specifically, of the 45% who confronted, 9% responded to 3 of 3 sexist comments, 11% responded to 2 of 3, and 25% responded to only 1 out of 3 sexist comments. Rounding these numbers up, and since other students had reported fewer objections, in our model we set 10% of agents to confront as targets with *p*=1, 10% with *p*=.66, and 20% with *p*=.33. These parameters are the same for male and female agents.

#### Confronting bias as an ally: Costs and likelihood

Czopp & Monteith (2003) found that allies are perceived less negatively after confronting biased behavior than targets. Participants’ perceptions of confronters’ overreaction as a function of whether the confrontation came from the target or a nontarget, were reported as follows: M(target) = 4.2 vs. M(nontarget) = 3.8 on a 7 point scale (Study 2). Said another way, confronting with target status relative to non-target/ally status increases negative reactions by .4 points on a 7 point scale (or 6%). Thus the normalized parameters indicate that if we set the cost of confrontation at 1 for women, the cost of confronting sexism should be .94 for men. Other studies have shown that third-party punishers are preferred over victims who punish unfair treatment (*28*). Taken together, we estimate a modest difference of .2 between the costs of target vs. ally objection, such that when a target (male or female) receives a cost of 1 for objection to bias, an ally (non-target gender) receives a cost of .8.

There are no estimates of the likelihood of ally confrontation in the literature (Benjamin J. Drury & Kaiser, 2014). However, confrontation assumes that the bias is perceived in the first place (Stangor, Swim, Van Allen, & Sechrist, 2002). According to a series of daily diary studies, college-aged women reported a median of 3.5 incidents of sexism directed at women per week whereas men reported 2 incidents of sexism directed at women per week (Swim, Hyers, Cohen, & Ferguson, 2001). Because we do not have confrontation base rates for men, we can estimate this rate as a function of gender differences in *perception* of bias. <u>We assume men and women are equally motivated to confront.</u> Therefore, if men perceive 57% of the sexist incidents women perceive, and 45% of women confront, we assume men confront in proportion: 26% of men confront when someone makes a sexist remark. (Future empirical work is required to determine the probability of ally objection and perception separately, and find interventions that train for bias perception and objection respectively.) Estimating the probability of objection as an ally was more challenging. In the absence of direct empirical estimates, we combined the ratios above with the information that men perceived instances of sexism noticed by women about 56% of the time. Based on this finding, we estimated the probability of objection as an ally as follows: 10% object as ally with .66 probability, and 10% object as ally with probability .33.

#### Does bias confrontation change subsequent likelihood of bias?

The data tentatively suggest yes. For example, men who were confronted by female confederates about making a sexist comment immediately engaged in compensatory efforts and later reduced their use of sexist language (Mallett & Wagner, 2011). A more recent study tested whether these effects of interpersonal confrontation on reducing prejudice endure (albeit in a racial bias context) and reported that effects of bias confrontation on prejudice were still present a week later (Chaney & Sanchez, 2018).

A study conducted a test of confrontation via twitter by varying the identity of the confronter (actually a bot) between in-group (white male) and out-group (black male) and the number of Twitter followers each bot had. They found that white male Twitter users who were confronted by a high-follower white male significantly reduced their use of a racist slur in the days that followed (Munger, 2017). Based on this study, we estimated that receiving confrontation decreased the perpetuator’s probability of expressing bias by 5%.

There is very little work on the effects of *not* confronting so we had to estimate parameters based on evidence from one study in the context of gender bias and two studies in the context of racial bias. Rasinski, Geers, & Czopp (2013) found that among individuals who said that confronting sexism was important to them, not confronting a sexist comment (relative to being exposed to a sexist comment without the opportunity to confront) decreased self-reported importance of confronting on a second measure. Specifically, not confronting decreased participants’ beliefs in the importance of confronting from 6.7 to 5.9 (estimated from figure): a difference of 0.8 on a 7-point scale. Of course we cannot assume belief change translates directly into behavior change. Thus we also looked at two studies (in the domain of racial bias confrontation) by Szekeres et al. (under review) who report a 9% and 12% decrease in out-group support (i.e., donation to Black student education program) after not confronting (relative to controls).

### Model implementation

#### What does each simulation look like?

We simulated environments with 100 agents that underwent 1000 “meetings” or conversational rounds in which 2-8 agents were sampled from the 100 population. The agents in each meeting/conversational round interacted and had the opportunity to display sexist behavior and then to objected to one another according to their probabilities and ratios.

#### Agents

We initialized each simulation environment with 100 agents that were either male or female, with a range of female to male gender ratio, and equal distributions of bias between the female and the male groups. Specifically, 50% of either gender population made zero biased comments, 10% of each group’s population has a .2 probability of bias, 10% has a probability of .4, 10% of .6, 10% .8, and 10% a probability of 1 of exhibiting gender bias in an interaction. Each agent was initialized with a probability of objecting to bias (see section above) as follows: 40% of female agents (10% with probability 1, 10% with probability .66, and 20% with probability of .33), and 20% of male agents (10% with probability .66 and 10% with probability .33) were likely to object overall.

#### Meetings

Each simulation consisted of a series of 1000 sequential “meetings”, in which 2-8 random agents were sampled from the population to interact. During each meeting, bias probabilities determined whether gender bias occured, whether objections took place, and then the agents’ probabilities of showing bias were updated based on their learning parameters. Receiving bias, confronting bias, and receiving confrontation incurred costs to the agents. Notably, agents could “learn” from the occurrence of bias and confrontation, such that their probability of future bias could increase (bias propagation) or decrease (bias inhibition).

**Supplementary Figure 1.**
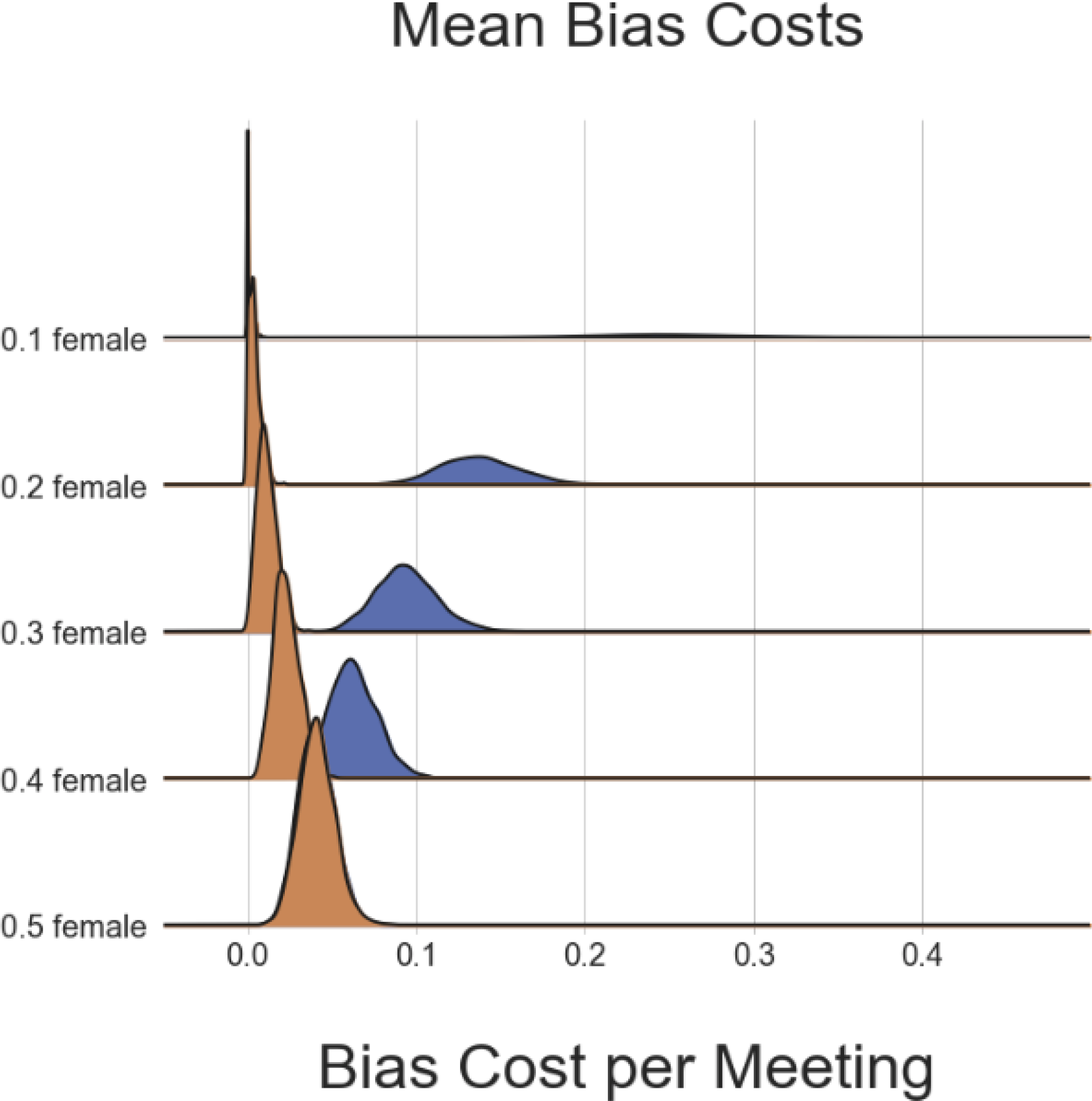
Mean bias cost for female (blue) and male (orange) agents, given no objections. Mean cost of bias.

**Supplementary Figure 2.**
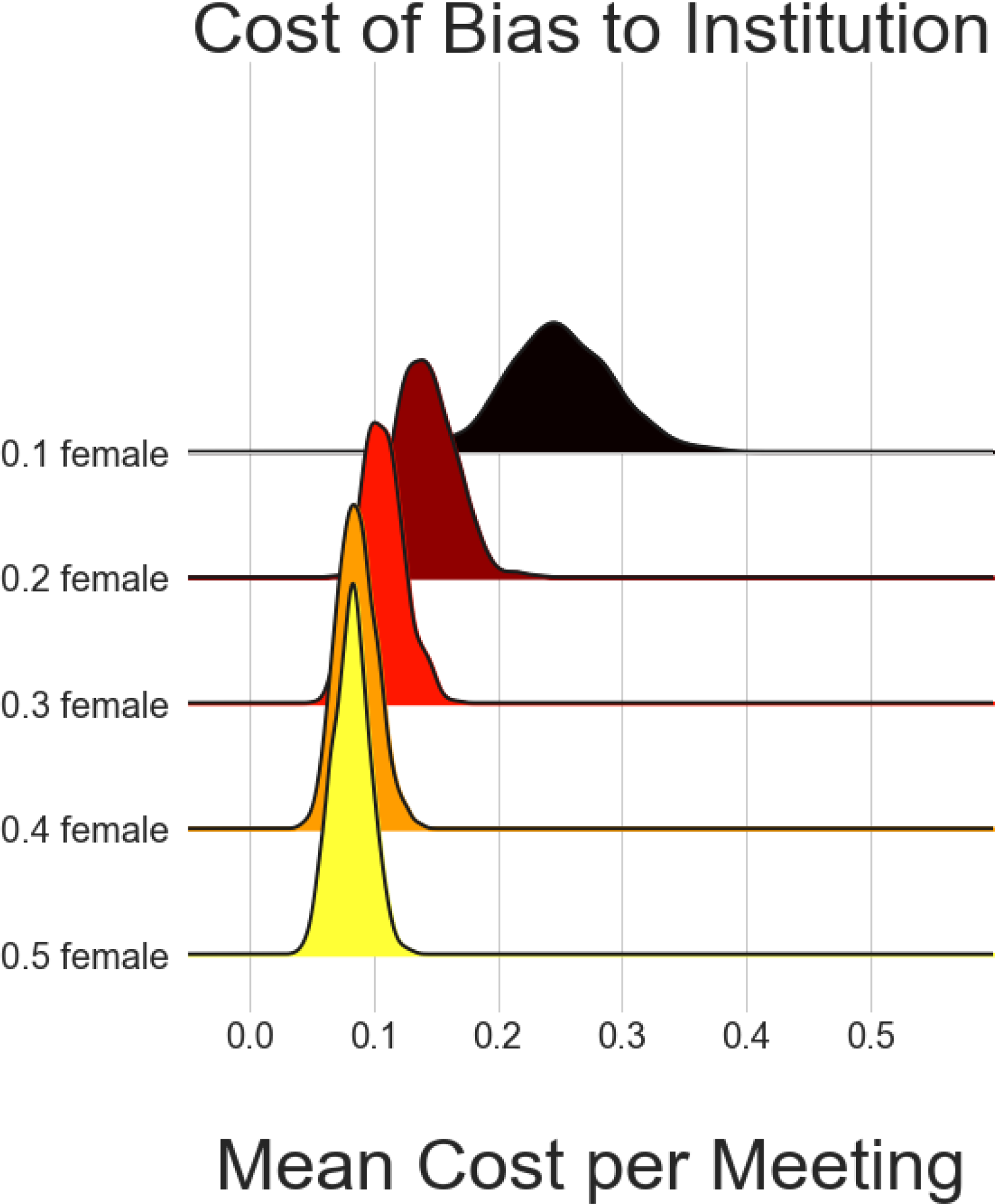
Mean bias cost to institutions, given no objections. Mean cost of bias.

**Supplementary Figure 3.**
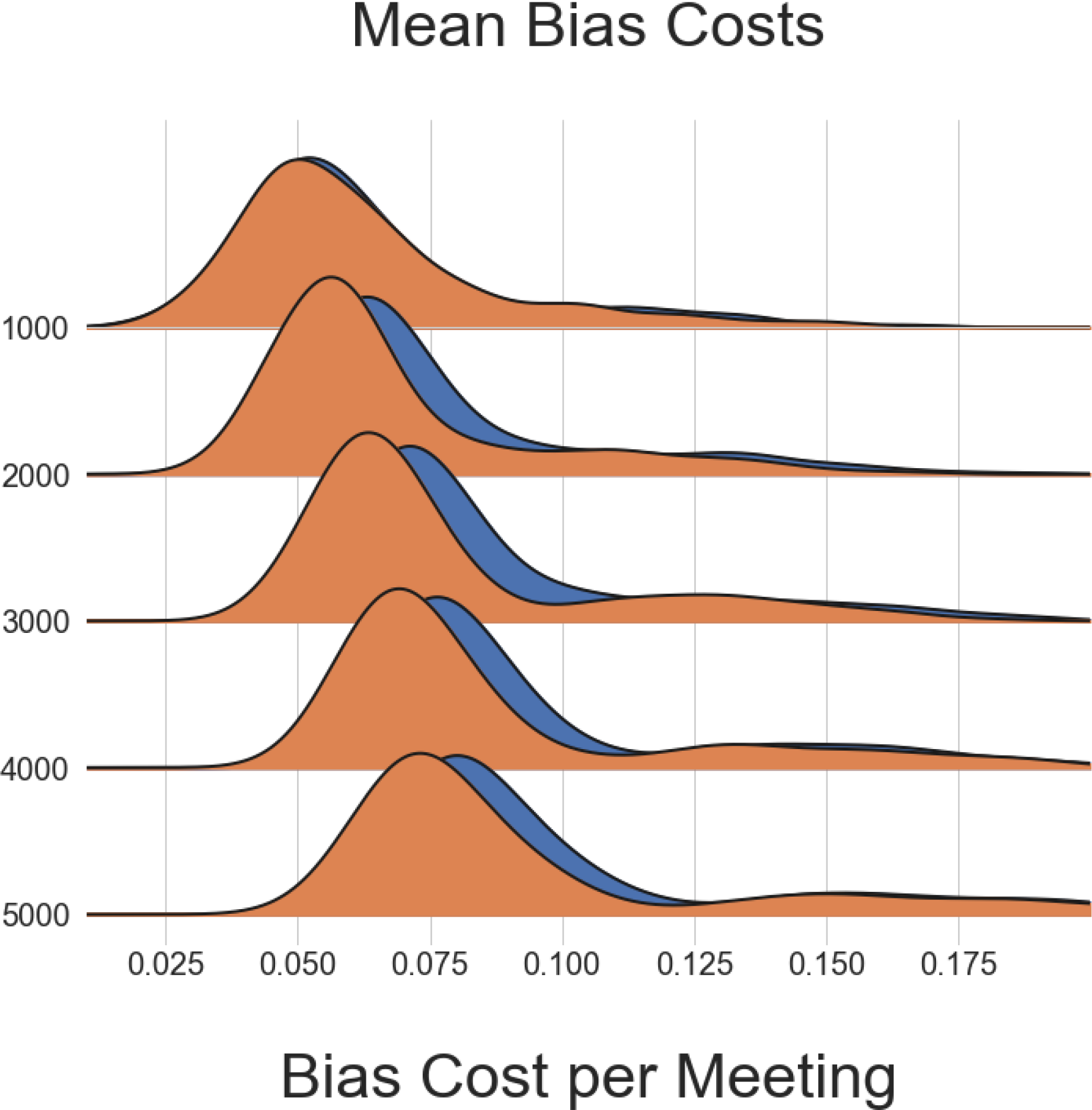
Cost of bias after equality policy. Y Axis: number of meetings.

